# ShcA promotes chondrocyte hypertrophic commitment and osteoarthritis in mice through RunX2 nuclear translocation and YAP1 inactivation

**DOI:** 10.1101/2020.07.16.206870

**Authors:** A. Abou-Jaoude, M. Courtes, L. Badique, D. Elhaj Mahmoud, C. Abboud, M. Mlih, H. Justiniano, A. Lemle, S. Awan, J. Terrand, A. Niemeier, A. Barbero, X. Houard, P. Boucher, RL Matz

## Abstract

Chondrocyte hypertrophic differentiation, a key process in endochondral ossification (EO), is also a feature of osteoarthritis leading to articular cartilage destruction. ShcA (Src homology and Collagen A) is an adaptor protein that binds to the cytoplasmic tail of receptor tyrosine kinases. We found that deletion of ShcA in chondrocytes of mice inhibits hypertrophic differentiation, alters the EO process, and leads to dwarfism. ShcA promotes ERK1/2 activation, nuclear translocation of the master transcription factor for chondrocyte hypertrophy, RunX2, while maintaining the Runx2 inhibitor YAP1 in its cytosolic inactive form. This leads to hypertrophic commitment and expression of markers of hypertrophy, such as Collagen X. In addition, ShcA deletion in chondrocytes protects from age-related osteoarthritis development in mice. Our results reveal that ShcA integrates multiple stimuli which affect the intracellular signaling processes leading to the hypertrophic commitment of chondrocytes and osteoarthritis.

## INTRODUCTION

Chondrocyte differentiation and hypertrophy are key events in long bones and vertebral skeleton formation allowing skeletal growth from embryogenesis to skeletal maturity (1). During endochondral ossification (EO), chondrocytes produce transient cartilage scaffolds for new bone formation in the growth plate (1). The growth plate is a highly organized cartilage structure in which chondrocytes proliferate and differentiate in pre-hypertrophic and then hypertrophic chondrocytes. Hypertrophic chondrocytes orchestrate cartilage extracellular matrix (ECM) remodeling, its calcification and osteoblasts infiltration which lead to replacement of cartilage by bone and longitudinal bone growth. Defective hypertrophic differentiation and ossification can lead to impaired longitudinal growth and thus dwarfism (2).

Chondrocyte hypertrophy is also a feature of osteoarthritis (OA) as quiescent articular chondrocytes can undergo an aberrant terminal hypertrophic differentiation (3, 4). The switch from a quiescent to a hypertrophic phenotype is accompanied by the pathologic remodeling of the ECM leading to articular cartilage destruction.

In the articular cartilage, and in the growth plate, resting chondrocytes, synthesize an ECM rich in collagen type II (col2a1) and proteoglycans. In addition to changes in cell morphology, collagen type II expression decreases during chondrocyte maturation, and the hypertrophic chondrocyte initiates the synthesis of collagen type X, together with proteolytic enzymes such as matrix metalloproteinase 13 (MMP13) which damage ECM integrity and lead to cartilage destruction (1, 3, 5).

Although the mechanisms involved are not fully understood, multiple factors, including matrix proteins, growth factors like IGF-I or FGF, transcription factors, and intracellular signaling proteins have been involved in chondrocyte hypertrophy (2)(6–8). Among intracellular signaling pathways, the MAPK/ERK1/2 pathway can be activated by various stimuli including growth factors, and is involved in chondrocyte differentiation from the pre-hypertrophic stage to the late hypertrophic stage during EO (9). Furthermore, the MAPK/ERK1/2 pathway phosphorylates and activates RunX2 (Cbfa1), a master transcription factor for chondrocyte hypertrophy and an indispensable collagen X transactivator (10) (11, 12) (13).

ShcA (Src Homology and Collagen A) is a cytosolic adaptor protein that binds to the cytoplasmic tail of growth factor receptors once activated, such as the IGF-I-receptor, the FGF receptor-3 and integrins (14) (6). ShcA recruitment to the plasma membrane leads to the activation of the Ras:Raf:MEK1:ERK1/2 pathway. ShcA is expressed in hypertrophic chondrocytes but its precise role during chondrogenesis remain unknown (15). As ShcA has the potential to signal downstream of several plasma membrane receptors involved in chondrogenesis, and chondrocyte hypertrophic differentiation, and upstream of the MEK/ERK1/2 pathway, we hypothesized that it might act as a checkpoint in chondrocyte differentiation. To study the role of ShcA in chondrocyte differentiation, we suppressed its expression specifically in chondrocytes using the Cre/lox technology in mice.

## METHODS

### Mice

All animal experimentations and procedures were approved by the Institutional Animal Care and Use Committee (IACUC) of the University of Strasbourg, France, and performed conform to the guidelines from Directive 2010/63/EU of the European Parliament on the protection of animals used for scientific purposes (authorization APAFIS#15477). C57/B6 mice carrying a ShcA allele into which loxP sites are integrated have been generated by gene targeting in embryonic stem cells. LoxP sites have been introduced upstream of exon 2 and downstream of exon 7 (ShcA^flox/flox^) (16). Cre-mediated recombination resulted in deletion of a 2-kb fragment containing the sequence encoding the PTB domain required for binding to phosphorylated receptors and for signaling activity. Chondrocyte specific p66, p52 and p46 ShcA inactivation was achieved by crossing transgenic mice carrying the Twist2-Cre transgene (The Jackson laboratory) with ShcA^flox/flox^ mice. Genotyping of the wild type (TwShcA+) and ShcA mutant (TwShcA-) mice by polymerase chain reaction (PCR) was performed as described using primers specific for ShcA (primers available upon request) (17). Animals were maintained on a 12-h light/12-h dark cycle. For *in vivo* analysis, one month old mice, three months old mice or two years old mice were used. For chondrocytes isolation, 8 to 10 days old mice were used. The agents used for euthanasia were ketamine (750 mg/kg) and xylazine (50 mg/kg), intraperitoneally.

### Chondrocyte isolation and culture

The costal cartilage as well as femoral and tibial articular cartilage were isolated from TwShcA+ and TwShcA- mice 7 to 10 days after birth and primary murine chondrocytes from hyaline cartilage were extracted and cultured as previously described (18).

### Cells culture and differentiation

The mouse chondroprogenitor cell line ATDC5 was purchased from Sigma-Aldrich, maintained in DMEM/F12 medium supplemented with 5% FBS and 2mM L-Glutamine and incubated in a humidified atmosphere containing 5% CO_2_ at 37°C. The primary murine chondrocytes were used to perform a three dimensional pellet culture model of chondrogenic differentiation *in vitro* as previously described (20). Briefly, after *ex vivo* expansion and dedifferentiation (5 passages), 500 000 cells were centrifuged in polypropylene tubes at 500g for 5 minutes. They were incubated in a humidified atmosphere containing 5% CO_2_ at 37°C in a chondrogenic medium containing TGF beta-1 (10 ng/ml) in DMEM medium supplemented with 10% FBS and 2 mM L-glutamine for up to 2 weeks. Medium was changed every 2-3 days. The primary murine chondrocytes were also used to perform a two dimensional culture model of hypertrophic differentiation *in vitro* as previously described (19). Briefly, after *ex vivo* expansion and dedifferentiation (5 passages), 300 000 cells/well were seeded in 6 well plates. The day after seeding, they were incubated in a humidified atmosphere containing 5% CO_2_ at 37°C in a chondrogenic medium containing TGF beta-1 (10 ng/ml) (Sigma Aldrich) in DMEM medium supplemented with 10% FBS and 2 mM L-glutamine for up to 2 weeks. Medium was changed every 2-3 days.

### Histology and immunostaining experiments

Mouse joints were isolated and fixed in 4% buffered formaldehyde (Formalin) for 5 days (one or three month old mice) or 7 days (two years old mice), then decalcified in 10% (w/v) EDTA disodium (pH 7.4) for 10 days (one or three month old mice) or 21 days (two years old mice) at room temperature before being embedded in paraffin. Longitudinal joint sections at 5 μm thickness were processed for Safranin O and Fast Green staining, hematoxylin/eosin or immunohistochemical staining according to standard methods. For immunohistochemical analysis the following antibodies were used: rabbit anti-collagen II (AbCam), rabbit anti-collagen X (AbCam), rabbit anti phospho-ERK1/2 (Cell signaling Technology). The Vectastain kit (Clinisciences) and the DAB detection system (Clinisciences) were used. The stained specimens were photographed digitally under a microscope.

For quantitative analysis of the hypertrophic zone of the growth plate, images taken through the microscope were processed using Image J^®^.

To evaluate osteoarthritis severity, two histopathology scorings were applied after Safranin O and Fast Green staining: the OsteoArthritis Research Society International (OARSI) and the modified Mankin scoring systems (20,21). The Mankin scoring system assigns grades to histological features characteristic of OA independently of the location or extent whereas the OARSI attributes stage to the horizontal extent and grades to the vertical depth within cartilage reflecting the aggressiveness of the lesions. OARSI scoring system was used in three sections with different depth. Sections were blinded and scored by three different experienced scientists. Averaged scores were used in statistical analyses. **Western blot -** SDS-polyacrylamide gel electrophoresis and immunoblot analysis were performed according to standard procedures. Proteins were transferred onto nitrocellulose membranes and immunoblot analyses were carried out using rabbit antibodies directed against ShcA (Millipore), Collagen X (Abcam), Collagen II (Abcam), MMP13 (AbCam), phospho ERK ½ (Cell Signaling Technology), RunX2 (Cell signaling), phospho-YAP1 (ser 127) and YAP1 (Cell Signaling Technology), GAPDH (Millipore). ImageQuant®LAS 4000 Imaging System (Amersham) was used to visualize protein expression. Optical densitometry was performed with Adobe Photoshop and Image J®.

### Cell fractionation

Cells were seeded in P100 dishes and, 24 hours after seeding, were transfected with either scrambled siRNA, or siRNA against p66, p52 and p46 isoforms of ShcA (Dharmacon) at a final concentration of 100 nM using lipofectamine 3000 (Thermo Fischer Scientific). 48 hours post-transfection, the cells were fractionated as previously described (22).

### Confocal Microscopy

Primary chondrocytes were seeded on glass slides, and 48 hours later were fixed with 3% paraformaldehyde, and incubated with anti-RunX2 (Cell signaling Technology), anti-YAP1 (Cell signaling Technology), anti-IgG control primary antibodies and Alexa Fluor 488 (Fischer Scientific) secondary antibodies. Immunofluorescence-labeled cells were analyzed using a Leica TSC SPE confocal microscope with the ×63 oil immersion objective.

### Statistical analysis

Values are reported as mean ± SEM of at least triplicate determinations. Statistical significance (*P* < 0.05) was determined using an unpaired Student’s *t* test (GraphPad Prism®, *Abacus Concepts, Berkeley, CA*). P-values < 0.05, < 0.01, <0.001 and < 0.0001 are identified with 1, 2, 3 or 4 asterisks, respectively. ns: p > 0.05.

## RESULTS

### Specific deletion of ShcA in chondrocytes leads to dwarfism

We generated TwShcA- mice in which ShcA is selectively ablated in chondrocytes by breeding Twist2 transgenic mice with ShcA^flox/flox^ mice (16). In TwShcA- mice, the three isoforms of ShcA were efficiently reduced in chondrocytes isolated from knee articular cartilage, vertebral and costal cartilage but not in non-cartilaginous tissues like lungs or the spleen (supplemental figure A and B). TwShcA- mice showed a dwarfism phenotype with a decrease in body size compared to control littermates (TwShcA+), (−25,4 +/- 1,3 %, p< 0,001) as well as in body weight (25,6 +/- 0,8 g TwShcA+ *versus* 19,1 +/- 0,9 g TwShcA-, p< 0,001), and tibial length (−30,6 +/- 1,4 %, p< 0,0001) at 12 weeks of age (figure 1A and D). Vertebral bodies and hind and front legs lengths were also significantly decreased at 12 weeks of age (figure 1A and B).

**FIGURE 1:**
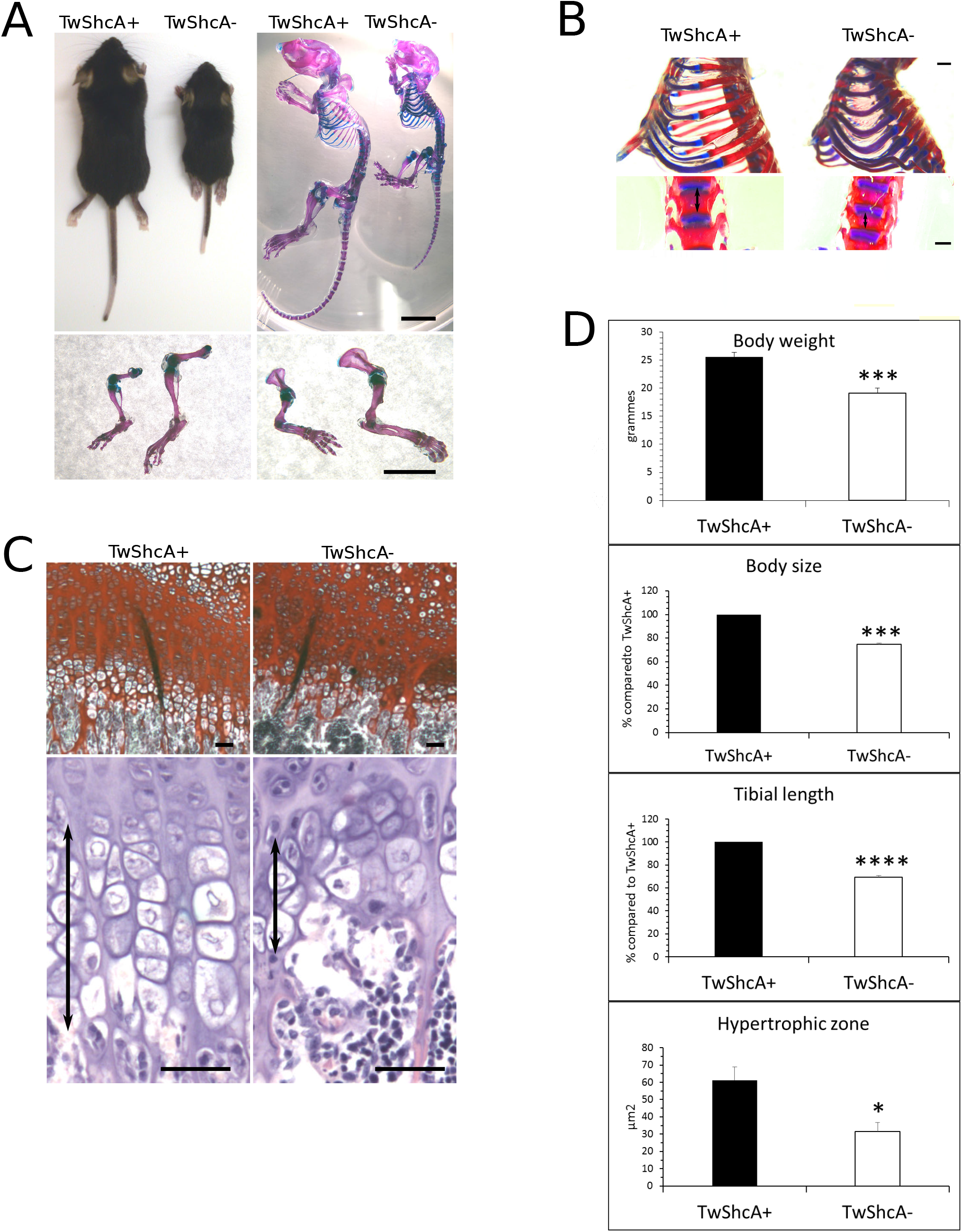
Dwarfism phenotype and decreased hypertrophic chondrocyte maturation in growth plate from TwShcA- mice. A) Representative body appearance and alizarin red- and alcian blue-stained bone and cartilage (upper panel) and alizarin red- and alcian blue-stained hind legs (lower left panel) and front legs (lower right panel) in mice that express (TwShcA+) or lack ShcA in chondrocytes (TwShcA-). Scale bars 10 mm. B) Alizarin red- and alcian blue-stained rib cage (upper panel) and spine (lower panel) from mice that express (TwShcA+) or lack ShcA (TwShcA-) in chondrocytes. White double headed arrow: alizarin-stained vertebral bone. Scale bars 1 mm. C) Safranin O-fast green (upper panel) and hematoxylin/ eosin (lower panel) stainings of tibial growth plate from mice that express (TwShcA+) or lack ShcA in chondrocytes (TwShcA-). Black double headed arrow: hypertrophic zone. Scale bars 100 μm. D) Quantification of body weight, size, tibial length and growth plate surface of mice that express (TwShcA+) or lack ShcA in chondrocytes (TwShcA-) (n= 7 mice in each group). * p< 0.05 *** p< 0.001 **** p>0.0001. Values are mean ± s.e.m. Two-tailed unpaired Student’s t-test.

Alizarin red staining for bone tissue and alcian blue staining for cartilage tissue showed an increased cartilage-to-bone ratio in the rib cage and the spine from TwShcA- mice compared to TwShcA+ mice (figure 1B). Histological analysis of safranin O- and hematoxylin-eosin-stained tibial growth plates indicated a disorganized, non-columnar proliferating chondrocytes zone and a shorter hypertrophic chondrocytes zone in one month old mice (61.14 ± 7,610 μm^2^ in TwShcA+ *versus* 31.54 ± 5.205 μm^2^ in TwShcA- mice, p<0,05) (Figure 1C and D). A similar decrease in the hypertrophic zone was observed in vertebral growth plates (supplemental figure C).

These data are indicative of an altered EO process and suggest that ShcA is required for chondrocyte terminal maturation towards hypertrophy.

### ShcA drives chondrocyte maturation to hypertrophy and collagen X expression

When primary chondrocytes isolated from hyaline cartilage are expanded in monolayer, they typically dedifferentiate and acquire a fibroblast-like phenotype (18). These fibroblast-like cells can be redifferentiated using chondrogenic differentiation protocols either in monolayer culture or in pellet culture (19). Using such protocols, we observed that, after dedifferentiation, chondrocytes isolated from TwShcA- mice are less prone to hypertrophic commitment than those isolated from TwShcA+ mice as shown by the decrease in the alizarin red stained mineralized matrix (Figure 2A). There are less hypertrophic chondrocytes in the pellet culture of chondrocytes isolated from TwShcA- mice compared to those from TwShcA+ mice (Figure 2B, safranin O staining). Also, the immunological staining of the pellets shows a decrease in collagen X staining, the main marker of hypertrophy, in chondrocytes isolated from TwShcA- mice compared to those from TwShcA+ mice (Figure 2B). These *in vitro* data indicate a role for ShcA in controlling chondrocyte hypertrophic commitment.

**FIGURE 2:**
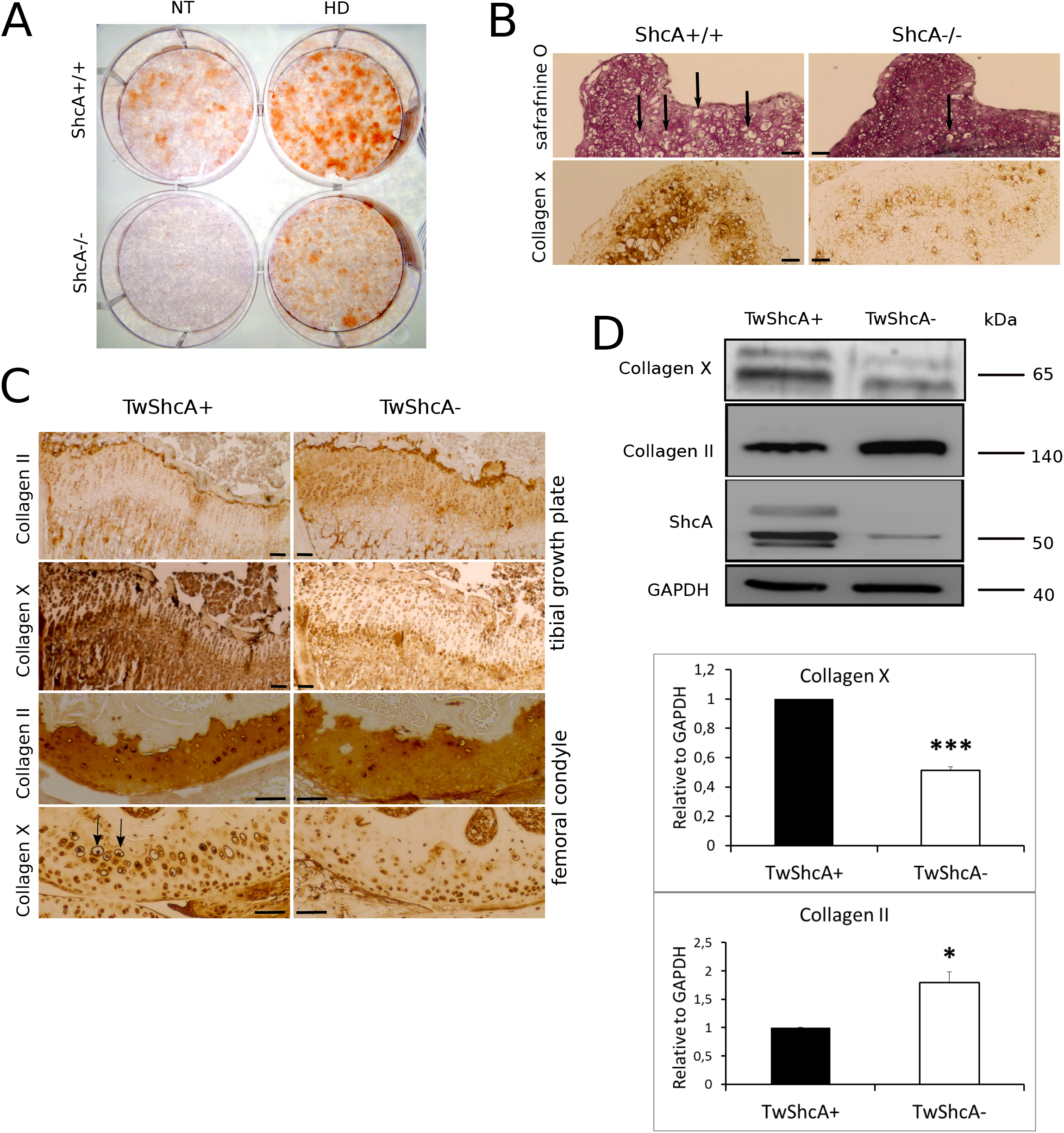
Decreased hypertrophic chondrocyte maturation and collagen X expression in TwShcA- mice and ShcA deficient cells. A) Alizarin red staining of articular chondrocytes isolated from mice that express (ShcA+/+) or lack ShcA in chondrocytes (ShcA-/-) submitted (HD) or not (NT) to an hypertrophic environment. B) Safranin O and collagen X staining of three dimensional pellet culture of articular chondrocytes isolated from mice that express (ShcA+/+) or lack ShcA in chondrocytes (ShcA-/-). Arrows show hypertrophic chondrocytes. Scale bars 100 μm. C) Collagen II and Collagen X staining of tibial growth plate (upper panels) and femoral condyle articular cartilage (lower panels) in mice that express (TwShcA+) or lack ShcA in chondrocytes (TwShcA-). Arrows show hypertrophic chondrocytes. Scale bars 100 μm. D) Western-blot analysis and relative quantification of collagen X and collagen II protein levels in knee joint articular chondrocytes isolated from mice that express (TwShcA+) or lack ShcA in chondrocytes (TwShcA-) (n= 7 in each group for collagen x, n=5 in each group for collagen II).

Collagen X is an ECM protein specifically synthetized by hypertrophic chondrocytes whereas collagen II is an ECM protein synthetized by quiescent chondrocytes (23, 24). Immuno-histological analysis of tibial growth plates sections from one month old mice indicated a marked decrease in collagen X staining as well as an increase in collagen II staining in TwShcA- mice compared to TwShcA+ mice (Figure 2C). In femoral articular cartilage from one year old mice, immunostaining of collagen X and the number of hypertrophic chondrocytes were decreased whereas immunostaining of collagen II was increased in TwShcA- mice compared to TwShcA+ mice (Figure 2C). Quantifications of collagen X and collagen II expressions show that collagen X was decreased by 40% in chondrocytes isolated from TwShcA- mice knee joint cartilage (1 *versus* 0.6 ± 0.05, TwShcA+ *versus* TwShcA- mice, p<0.0001) whereas collagen II expression was significantly increased (1 *versus* 1.8 ± 0.19, in TwShcA+ *versus* TwShcA- mice, p<0.01) (Figure 2D).

Not all hypertrophic markers were downregulated in ShcA-deficient chondrocytes as the expression of MMP13 was not significantly different between chondrocytes from TwShcA+ and TwShcA- mice (1 *versus* 1.2 ± 0.24, NS) (Supplemental figure D). The matrix protease MMP13 is a late hypertrophic marker (25), suggesting that ShcA is involved in the earlier stages of hypertrophic differentiation.

Thus, the decrease of the hypertrophic marker collagen X parallels the inhibition of chondrocyte hypertrophic commitment in the absence of ShcA.

### ShcA induces hypertrophic commitment by promoting ERK1/2 activation, RunX2 nuclear translocation and by retaining YAP1 in its cytosolic inactive phosphorylated form

One of the main downstream target of ShcA is ERK1/2 (14). It has been reported that the MAPK/ERK1/2 pathway promotes chondrocytes differentiation from the pre-hypertrophic to the late hypertrophic stage during endochondral ossification (9). Once activated ERK1/2 phosphorylates and activates Runx2 in osteoblasts (10). Runx2 and its target gene collagen X are essential for chondrocyte hypertrophy (11, 13). Thus, by activating ERK1/2, ShcA might induce RunX2 activation leading to Collagen X expression and chondrocyte hypertrophic commitment.

To test this, we first quantified ERK1/2 phosphorylation in primary chondrocytes isolated from knee joint cartilage. We found that the deletion of ShcA leads to a marked decrease in p-ERK1/2 compared to controls (1 *versus* 0.51 ± 0.06, p<0.01) (Figure 3A). A 50% decrease in the expression of phospho-ERK1/2 was also observed *in vivo* in tibial growth plate hypertrophic chondrocytes from one month old TwShcA- mice (Figure 3B). In columnar proliferating chondrocytes, the decrease was observed to a lesser extent (Figure 3B).

**FIGURE 3:**
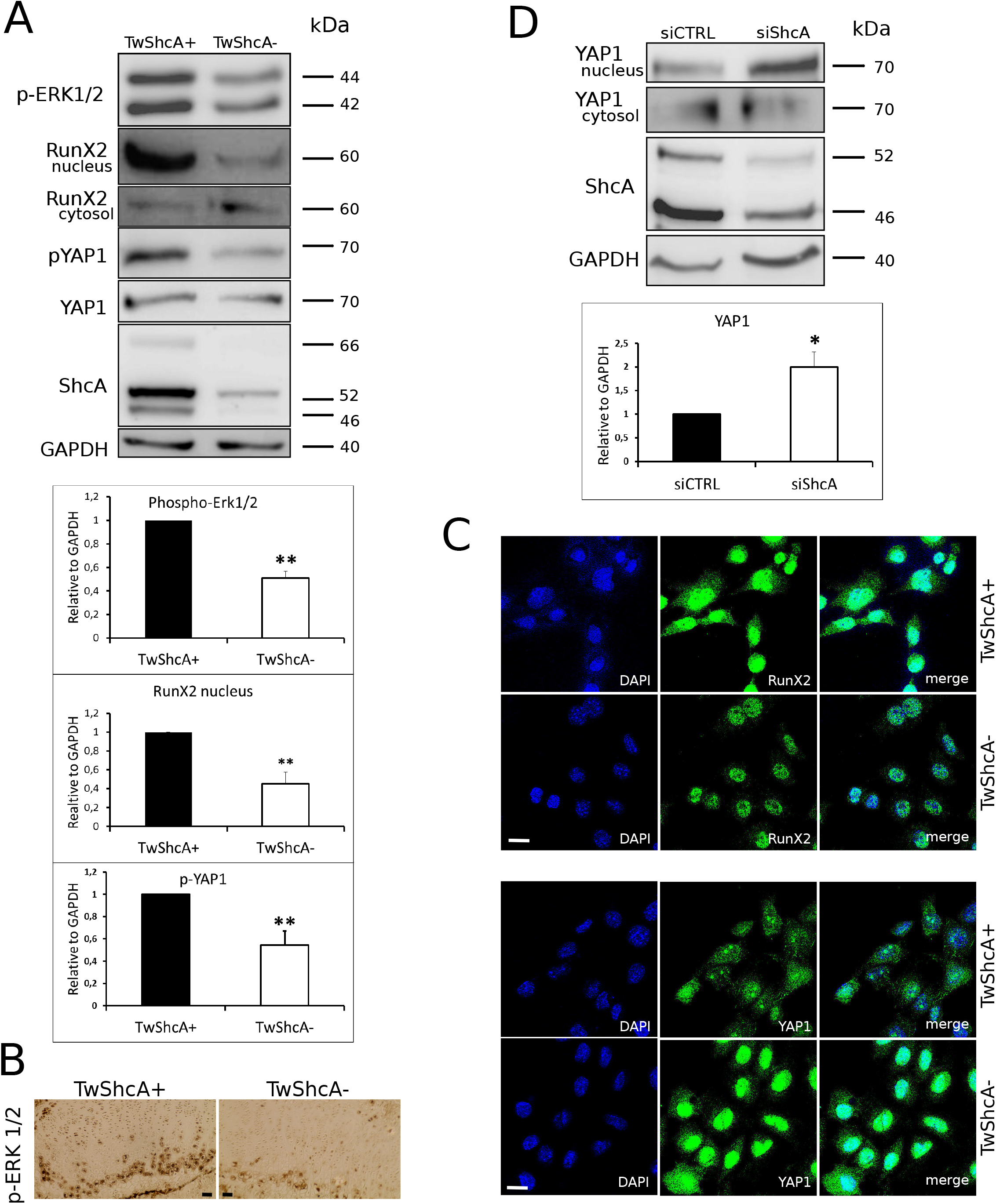
Decreased ERKl/2 and RunX2 activation and increased YAP1 activation in TwShcA- mice and ShcA deficient cells. A) Western blot analysis and relative quantification of phospho-ERK1/2, nuclear and cytosolic RunX2, phospho-YAP1, YAP1, ShcA and GAPDH proteins levels in knee joint articular chondrocytes isolated from mice that express (TwShcA+) or lack ShcA in chondrocytes (TwShcA-) (n= 8 mice in each group for phospho-ERK1/2 ShcA, n= 5 mice in each group for RunX2, phospho-YAP1, YAP1). B) phospho-ERK1/2 staining of tibial growth plate in mice that express (TwShcA+) or lack ShcA in chondrocytes (TwShcA-). Scale bars 100 μm. C) Representative confocal immunostaining of RunX2 (upper panel) and YAP1 (lower panel) in articular chondrocytes isolated from mice that express (TwShcA+) or lack ShcA in chondrocytes (TwShcA-) (n= 3 separate experiments). Scale bars 10 μm. D) Western-blot analysis and relative quantification of nuclear and cytosolic YAP1 protein levels in ATDC5 cells down-regulated for ShcA (siShcA) and control cells (siCTRL) (n= 4 experiments in each group). *p<0.05, **p<0.001. Values are mean ± s.e.m. Two-tailed unpaired Student’s t-test.

We next tested whether ShcA promotes Runx2 nuclear translocation. Using cell fractionation and immuno-fluorescence experiments in primary chondrocytes isolated from knee joint cartilage of TwShcA+ and TwShcA- mice, we found a marked decrease in RunX2 expression in the nucleus in ShcA deficient cells (1 *versus* 0.45±0.12, p<0.01, cell fractionation) (Figure 3A) and a decreased nuclear staining of RunX2 in ShcA deficient cells (Figure 3C, confocal microscopy). This indicate that ShcA is required for RunX2 nuclear translocation in chondrocytes.

YAP1 is a transcriptional effector of the Hippo pathway. In cells, YAP1 is present in a cytosolic Ser/Thr phosphorylated inactive form (p-YAP1), whereas in the nucleus YAP1 regulate transcription (27). Because YAP1 can bind to RunX2 and suppresses collagen X transcription (26, 27), we tested whether ShcA retains p-YAP1 in the cytosol and thus prevents its nuclear translocation. Using primary chondrocytes isolated from knee joint cartilage of TwShcA+ and TwShcA- mice, we found a significant decrease in p-YAP1 expression in ShcA deficient cells (1 *versus* 0.54±0.12, p<0.01) (Figure 3A). We also tested YAP1 activation by its nuclear translocation in presence or absence of ShcA. Using cell fractionation and immuno-fluorescence experiments we found a marked increase in YAP1 nuclear translocation in ShcA deficient cells (1 *versus* 2.0±0.31, p<0.05) (Figure 3D) and a marked staining of YAP1 in the nucleus of ShcA deficient cells (Figure 3C).

Taken together, our results show that ShcA controls hypertrophic differentiation and collagen X expression by promoting ERK1/2 activation and RunX2 nuclear translocation, and by retaining YAP1 in its cytosolic inactive phosphorylated form.

### ShcA deletion in chondrocytes protects from aged-related OA development in mice

Aberrant terminal hypertrophic differentiation of articular chondrocytes has been implicated as a crucial step in OA pathogenesis (3, 4). During OA, articular chondrocytes change their phenotype to one resembling hypertrophic growth plate chondrocytes and OA can be regarded as an ectopic recapitulation of the endochondral ossification process (3, 4, 28).

Because ShcA promotes hypertrophic commitment, we tested whether its deletion protects against OA in aged TwShcA- mice compared to young TwShcA- mice. As mice of the C57BL/6 background are characterized by a determined propensity to develop spontaneous OA with age (29, 30), the TwShcA- mice were backcrossed on a C57BL/6 genetic background.

We then characterized the effect of ShcA deletion on spontaneous aged-induced OA development. Safranin O fast green staining of tibio-femoral joints showed a slightly increased glycosaminoglycan staining of tibial plateau and femoral condyle in one year old TwshcA-mice compared to TwShcA+ mice (Figure 4A). With aging, knee joints from TwShcA+ mice demonstrated erosion with loss of articular cartilage tissue staining, including in superficial and in at least portions of deeper cartilage layers, denudation, with matrix loss extending to calcified cartilage interface, and clefts to calcified zone (Figure 4B). Cartilage histopathology scorings, according to the OARSI and the modified Mankin scoring systems, showed a drastic increase in two years old TwShcA+ mice compared to young mice (OARSI: 12.5 ± 2.24 *versus* 0, Modified Mankin: 7.0 ± 0.89 *versus* 0, aged *versus* young mice), which validated the age-related development of osteoarthritic lesions.

**FIGURE 4:**
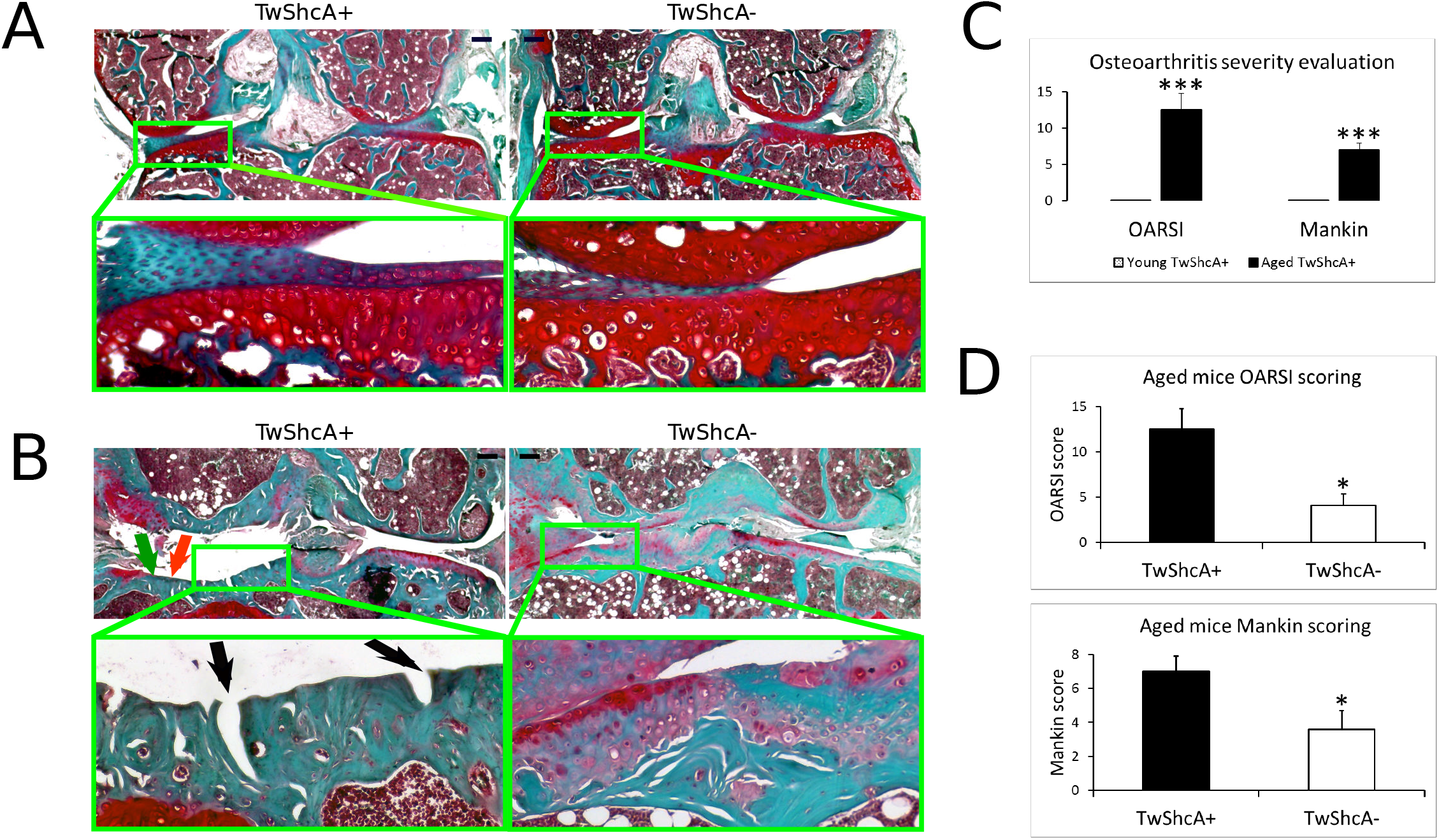
Inhibition of OA development in TwShcA- mice. Safranin O fast green staining of knee joint from one month old (A) and two years old (B) TwShcA+ and TwShcA- mice. Scale bars 250 μm (upper panels) 100 μm (lower panels). Green arrow: loss of articular cartilage, orange arrow: denudation of cartilage surface, black arrows: clefts to calcified zone. n= 5 mice in each group.

Cartilage histopathology scorings also showed a significant increase in aged TwShcA- mice compared to young TwShcA- mice, however the impairment of the cartilage tissue was substantially decreased compared to TwShcA+ mice at the same age (OARSI: 12.5 ± 2.2 *versus* 4.1 ± 1.3, p<0.05; modified Mankin: 7.0 ± 0.9 *versus* 3.6 ± 1.1, p<0.05, two years old TwShcA+ mice *versus* two years old TwShcA- mice).

These data indicate that ShcA promotes aged-related cartilage destruction and that deletion of ShcA in chondrocytes can slow down OA development in mice.

## SUPPLEMENTAL FIGURE

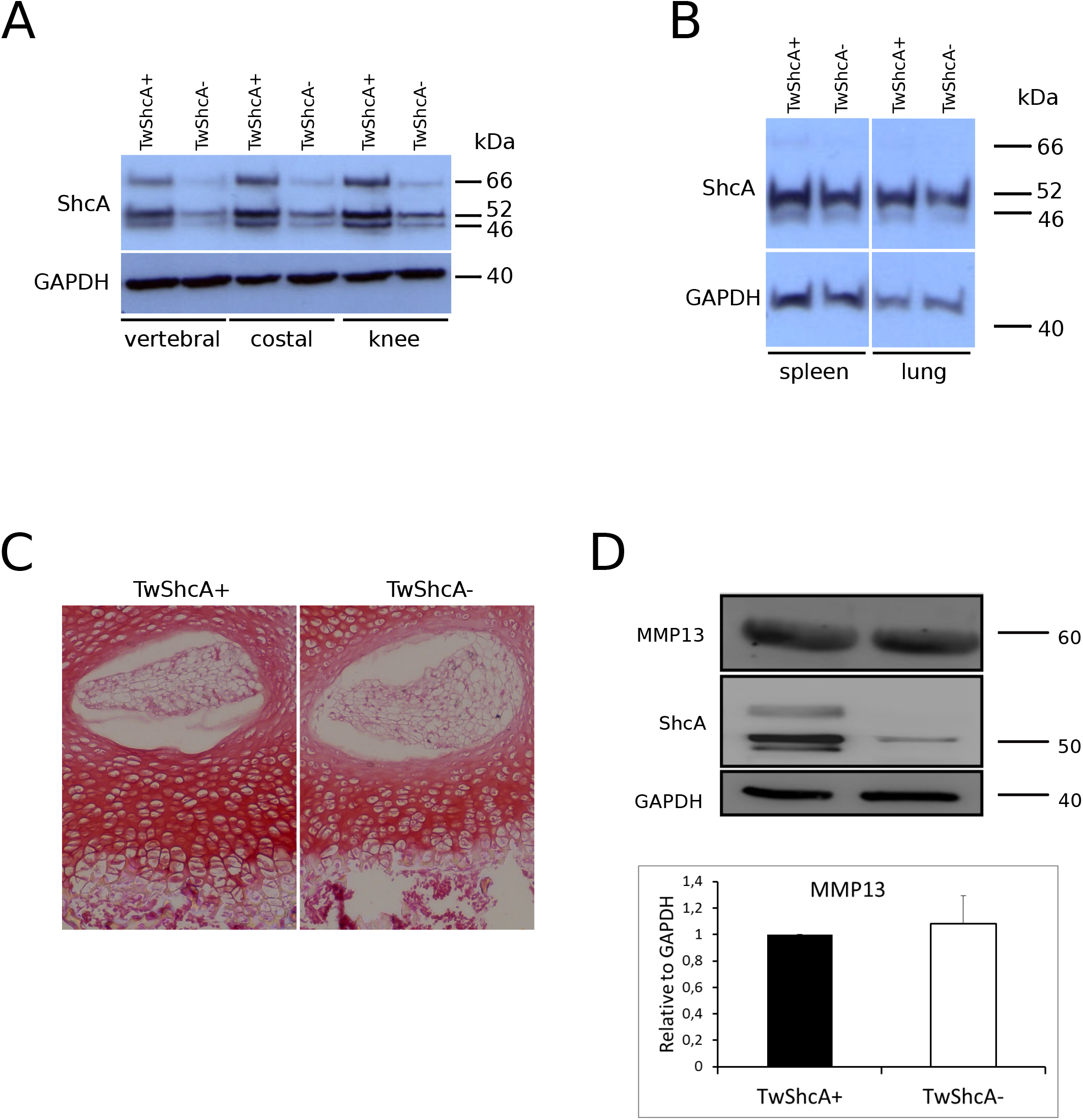
A) Expression of ShcA in vertebral, costal and knee articular cartilage from mice that express (TwShcA+) or lack ShcA in chondrocytes (TwShcA-). Primary chondrocytes were isolated from cartilage tissue of 8 to 10 days old TwShc+ and TwShcA- mice. B) Expression of ShcA in the spleen and lungs from TwShcA and TwShcA mice. Tissues were isolated from 3 months old TwShcA+ and TwShcA- mice. C) Decrease in the hypertrophic zone surface of vertebral cartilage in TwShcA- mice compared to TwShcA+ mice. D) Western-blot analysis and relative quantification of MMP13 protein levels in knee joint articular chondrocytes isolated from mice that express (TwShcA+) or lack ShcA (TwShcA-) in chondrocytes (n=4 in each group for MMP13).

## DISCUSSION

Integrins and numerous growth factors receptors are involved in chondrogenesis and terminal hypertrophic chondrocyte differentiation (2, 6–8). ShcA is a ubiquitously expressed adaptor protein that binds to the cytoplasmic tail of integrins and growth factor receptors once activated (14). Subsequently ShcA recruits and activates the Grb2:Sos:Ras:Raf:MEK1/2:ERK1/2 signaling cascade (14). Even if ShcA has been identified in hypertrophic chondrocytes, its function in chondrocyte differentiation has never been addressed. By its potential to integrate multiple extracellular stimuli ShcA may behave as an important regulator of chondrocyte differentiation. Our data indicate that specific deletion of ShcA in chondrocytes leads to a reduced cartilage-to-bone ratio and a dwarfism in mice. This phenotype is characterized by an altered EO process with an important inhibition of chondrocyte hypertrophic maturation in the growth plate. *In vitro* experiments confirmed the crucial role of ShcA in promoting chondrocyte maturation to hypertrophy. ShcA promotes chondrocyte hypertrophic commitment and osteoarthritis through RunX2 activation and YAP1 inhibition.

During hypertrophic maturation, chondrocyte-synthetized ECM changes, and while collagen II synthesis is lost, the expression of collagen X is initiated, along with MMP13 synthesis creating a favorable environment for mineralization and replacement of cartilage by bone (1) (31) (32). In the absence of ShcA, we observed a decrease in collagen X expression both in the growth plate and the articular cartilage from adult mice, but no decrease in collagen II expression. Instead, the expression of collagen II was increased. Thus, not only are ShcA-deficient chondrocytes refrained from undergoing hypertrophic differentiation but also they exhibit the collagen II marker of quiescence.

ERK1/2 is one of the main downstream targets of ShcA (14). Conditional deletion of ERK1/2 in hypertrophic chondrocyte leads to a decrease in long bones growth after birth and to an inhibition of the transition of early hypertrophic chondrocytes to terminally differentiated chondrocytes (9). We found that upon ShcA knockdown, the phosphorylation of ERK1/2 is decreased in chondrocytes. Interestingly, the decrease in ERK1/2 phosphorylation is mainly observed in hypertrophic chondrocytes from the growth plate and to a lesser extent in columnar proliferating chondrocytes. It has been reported that the main role of ERK1/2 in cartilage is to stimulate not cell proliferation but rather chondrocyte maturation and hypertrophic differentiation, and that c-Raf may be responsible for ERK1/2 activation in hypertrophic chondrocytes (9, 33). Our results demonstrate that upstream of c-Raf, ShcA is necessary to activate ERK1/2, a determinant factor for hypertrophic differentiation.

Runx2 has been implicated as a master transcription factor for chondrocyte hypertrophy (11). After its nuclear translocation, RunX2 can be phosphorylated and activated by ERK1/2 leading to its binding to the Collagen X promoter and transcriptional activation (10, 13, 34). We found that the decreased activation of ERK1/2 in ShcA-deficient chondrocytes correlates with a decreased nuclear translocation of RunX2. Taken together these observations suggest that ShcA activates chondrocyte hypertrophic differentiation and collagen X expression by activating ERK1/2 and by promoting RunX2 nuclear translocation. We cannot rule out the participation of other signaling pathways in RunX2 activation. Indeed, the mammalian Ste20-like kinase (MST) pathway or Hippo pathway was reported to inhibit RunX2 activation by phosphorylation of its serine 339 and 370 residues (35). And MST 1/2 kinases are negatively regulated by c-Raf (36). Hence, upstream of Raf, ShcA might also activate RunX2 by controlling a c-Raf-MST1/2 pathway.

Our study also reveals that not only ShcA drives chondrocyte maturation to hypertrophy by positively activating ERK1/2 and RunX2 but also by negatively regulating YAP1. We report that upon ShcA knockdown, the cytoplasmic inactive form of YAP1 is decreased and YAP1 nuclear translocation is increased. It has been shown that YAP1 can inhibit collagen X expression by a direct interaction with RunX2 (26, 27).

Our data highlight the crucial role of ShcA in regulating the nuclear access of the transcription factor RunX2 and its regulator YAP1 to control protein expression. In chondrocytes, we found that ShcA retains YAP1 in its inactive form in the cytoplasm while promoting ERK1/2 activation and RunX2 nuclear translocation. RunX2 nuclear translocation activates hypertrophic commitment and collagen X transcription. The ShcA-mediated retention of YAP1 in the cytoplasm might involve the formation of a ShcA-Grb2-YAP1 complex. Indeed, it has been described that YAP1 is able to interact with SH3 domain-containing proteins through its WW domain (37). Grb2 contains such a SH3 domain and is able to bind ShcA through its SH2 domain (38, 39).

Aberrant terminal hypertrophic differentiation of articular chondrocytes has been implicated as a crucial step in OA pathogenesis (3, 4). During this switch, articular chondrocytes change their phenotype to one resembling hypertrophic growth plate chondrocytes and OA can be regarded as an ectopic recapitulation of the endochondral ossification process (3, 4, 28). Our results show that the ShcA-controlled hypertrophic differentiation is also a mechanism involved in aged-related osteoarthritis development in mice. Chondrocyte specific ShcA-deficient mice are refrained from severe osteoarthritis.

Several initial events are involved in chondrocyte differentiation towards hypertrophy, i.e. mechanical stimuli through integrins or DDR2, growth factors receptors or LRP5/6 activation (40, 41). ShcA potentially binds to the cytoplasmic tail of these receptors (42) (14)(6). Our results reveal that ShcA behaves as a major regulator to integrate multiple stimuli and to complete the whole intracellular signaling process leading to hypertrophic commitment either in physiological processes like skeletal growth or in a pathological process like OA. By its potential to lock chondrocytes in a desired differentiation stage and to stop inadvertent hypertrophic differentiation, ShcA might represent an interesting therapeutic target in OA and cartilage tissue engineering.

## ACKNOWLEDGEMENTS

We are grateful to Brigitte Pollet and Lionel Host (UMR CNRS 7021) for technical assistance. This study was supported by grants from Société Française de Rhumatologie.

